# RNAseq dataset describing transcriptional changes in cervical sensory ganglia after bilateral pyramidotomy and forelimb intramuscular gene therapy with AAV1 encoding human neurotrophin-3

**DOI:** 10.1101/415877

**Authors:** Claudia Kathe, Lawrence D F Moon

## Abstract

Unilateral or bilateral corticospinal tract injury in the pyramids of adult rats causes changes in proprioceptive axon terminal arborization in the cervical spinal cord accompanied by hyperreflexia and abnormal movements including spasms [1, 2]. Treatment of affected forelimb muscles with an Adeno-Associated Viral Vector (AAV) encoding human neurotrophin-3 (NT3) normalizes many of these anatomical, neurophysiological and behavioural changes [1]. Interestingly, in several studies, neurotrophin-3 protein accumulates in cervical dorsal root ganglia (DRG) on the side ipsilateral to AAV injection [1, 3]. We hypothesize that neurotrophin-3 induces these changes (in proprioceptive axon wiring, proprioceptive reflex neurophysiology and sensorimotor behaviors involving proprioception) by modifying gene expression in affected cervical dorsal root ganglia (DRG). As a first step in testing this hypothesis, we analyzed the transcriptomes of cervical DRGs obtained during a previous study from naïve rats and from rats after bilateral pyramidotomy (bPYX) with unilateral intramuscular injections of either AAV1-CMV-NT3 or AAV1-CMV-EGFP made 24h after injury [1]. Ten weeks after surgery, Poly(A) RNAs and small RNAs from C6 to C8 DRGs on the treated side were sequenced. We detected mRNAs or small RNAs that were significantly regulated under three conditions (bPYX+GFP vs naïve; bPYX+NT3 versus naïve; bPYX+NT3 vs bPYX+GFP). We identified mRNAs and small RNAs whose expression level was altered after pyramidotomy and normalized by neurotrophin-3 treatment. A bioinformatic analysis enabled us to identify genes that are likely to be expressed in proprioceptors after injury and which were regulated by neurotrophin-3 in the direction expected from other datasets involving knockout or overexpression of neurotrophin-3. This dataset will help us and others identify genes in sensory neurons whose expression levels are regulated by neurotrophin-3 treatment. This may help identify novel therapeutic targets to improve sensation and movement after neurological injury. Data has been deposited in the Gene Expression Omnibus (GSE82197).

## Specifications Table

|  |  |
| --- | --- |
| Subject area | *Biology* |
| More specific subject area | *Neuroscience* |
| Type of data | *RNA sequencing data, immunohistochemistry* |
| How data was acquired | *RNA sequencing (Illumina HiSeq 2500)* |
| Data format | Analyzed |
| Experimental factors | Three groups of rats: 1) bilateral pyramidotomy plus intramuscular injection of AAV1-CMV-NT3; 2) bilateral pyramidotomy plus intramuscular injection of AAV1-CMV-EGFP; 3) naïve, unoperated rats |
| Experimental features | Analysis of differential expression of poly A RNAs and small RNAs from cervical dorsal root ganglia of three groups of rats 10 weeks after surgery |
| Data source location | London, UK |
| Data accessibility | The data will be available with this article. All RNAseq data are available in the GEO archive under the accession number GSE82197<br>http://www.ncbi.nlm.nih.gov/geo/query/acc.cgi?token=avgpicgcjhknzyv&acc=GSE82197. |

## Value of the data

- Our data show that gene expression in sensory dorsal root ganglia was modified by supraspinal injury
- Our data show that expression levels of some genes in sensory dorsal root ganglia were normalized by intramuscular overexpression of neurotrophin-3
- Our data are valuable because one may seek the overlap between this and related datasets to find genes that are regulated in sensory dorsal root ganglia by neurotrophin-3: some of the gene expression levels changed in the same direction as that predicted by work by other groups.
- Therapeutic modification of gene expression in sensory ganglia may lead to functional consequences.

## Data

The data presented here are related to the research article entitled “Intramuscular Neurotrophin-3 normalizes low threshold spinal reflexes, reduces spasms and improves mobility after bilateral corticospinal tract injury in rats “[1]. We now present additional data that were gathered as part of that work. Adult rats received bilateral transection of the corticospinal tracts in the medullary pyramids (bPYX). Twenty-four hours later, rats received unilateral injection into their forelimb flexors of either an Adeno-associated viral vector (AAV1) encoding human neurotrophin-3 (NT3) or Green Fluorescent Protein (GFP). These rats underwent behavioural testing, neurophysiological assessment and nerve tracing and tissues were recovered for histology as described elsewhere [1]. Owing to the facts that we found evidence that intramuscular neurotrophin-3 affects spinal networks via proprioceptive afferents, we decided to investigate transcriptional changes in the ipsilateral cervical DRGs. C6-C8 cervical dorsal root ganglia (DRG) from the treated side were removed ten weeks after transection. DRG were pooled from the treated side, homogenized and Poly(A) RNAs and small RNAs were sequenced separately.

RNA sequencing of poly-A RNAs revealed genes that were expressed in cervical DRG of unoperated (naïve) adult rats (Supplementary Table 2). RNA sequencing also revealed small RNAs that were expressed in these cervical DRG (Supplementary Table 3). The most abundant mRNAs in these DRG included microtubule-associated proteins (Map 1b, NM_019217; Map 1a, NM_030995), neurofilaments (Nefl, NM_031783; Nefm NM_017029; Nefh NM_012607) and Nogo-A (Rtn4, NM_031831). Other genes that were expressed at lower levels included markers of neurons of different sizes including TrkA (Ntrk1, NM_021589) and two isoforms of TrkB (Ntrk2, NM_012731 and NM_001163168). Markers of large DRG neurons were detected including Nefh, Parvalbumin (NM_022499) and Spp1 (osteopontin; NM_012881) [4, 5]. Markers of non-neuronal cells were also detected including satellite glia (GFAP NM_017009, or glutamine-ammonia ligase (glutamine synthetase NM_017073), markers of endothelial cells (CD31/PECAM1 NM_031591 [6]) and markers of inflammatory cells (Cd3g NM_001077646; CD163 NM_001107887).

Neurotrophin-3 signals through its canonical receptor TrkC which is known to be expressed in many large neurons as shown by immunolabelling, *in situ* hybridization and RNAseq of purified subpopulations of DRG neurons [4, 7, 8]. In our experiment, no isoform of TrkC (Ntrk3, NM_019248, NM_182809.2 or NM_001270656) was detected, probably because our samples consisted of homogenized DRG in which TrkC expressing cells form only a small proportion of all neurons [∼10% of cells in lumbar DRG are neurons [9] and ∼17% of neurons are TrkC positive in L4/5 DRG [8] thus the overall proportion of TrkC expressing cells will be ∼1.7% of cells in DRG].

Bilateral pyramidotomy plus intramuscular injection of AAV1-CMV-EGFP (bPYX+GFP) caused regulation of a small number of poly(A) RNAs, as assessed 10 weeks after surgery (relative to naïve rats; FDR<0.1; Supplementary Table 4) including downregulation of Ugt8 (a protein involved in myelin formation) and cyclin D1 (involved in the cell cycle and apoptosis). With looser stringency (p<0.05), 298 genes appeared regulated (Supplementary Table 4); for example, Frizzled-related protein (Frzb) was upregulated. Frzb has been shown to be upregulated in DRG neurons following peripheral but not central branch injury [10] and during primary afferent collateral sprouting of uninjured DRG neurons [11]. Bilateral pyramidotomy plus intramuscular injection of AAV1-CMV-NT3 (bPYX+NT3) relative to naïve rats (FDR<0.1) caused regulation of a smaller number of poly(A) RNAs, assessed 10 weeks after surgery (Supplementary Table 4) including Ugt8 (as above). With looser stringency (p<0.05), 89 genes were regulated (Supplementary Table 4). Comparison of bPYX+NT3 versus bPYX+GFP revealed evidence for regulation of 163 genes (p<0.05) but a higher stringency analysis (FDR<0.1) revealed no genes.

Comparison of these three lower stringency lists (Supplementary Table 4) showed that forty-eight mRNAs and eighteen small RNAs were dysregulated by injury (p<0.05, bPYX+GFP vs naïve; Supplementary Table 6) and stayed dysregulated with neurotrophin-3 (p>0.05, bPYX+NT3 vs bPYX+GFP; Supplementary Table 6) indicating the NT3 was not able to reverse expression of these genes after pyramidotomy. Seven mRNAs and seven small RNAs were dysregulated by neurotrophin-3 (p<0.05, bPYX+NT3 vs bPYX+GFP or bPYX+NT3 vs naïve; Supplementary Table 8) and not by injury alone (bPYX+GFP vs naive; Supplementary Table 7) indicating that some genes were modified by NT3 and not by pyramidotomy alone.

We were most interested in genes in sensory neurons whose levels were dysregulated by bilateral pyramidotomy and that were also normalized with neurotrophin-3 treatment. Thirty-two mRNAs and nine small RNAs were significantly dysregulated after injury (bPYX+GFP vs naïve rats) and normalized with neurotrophin-3 treatment (p<0.05, bPYX+NT3 vs bPYX+GFP animals), *i.e.*, indistinguishable from naive levels (Figure 1, Figure 2, Supplementary Table 7). For example, *Gas7* and *Sprr1a*, cytoskeletal and growth cone modelling proteins, were both found downregulated after injury, but were normalized with neurotrophin-3 (Figure 3). These and other genes may mediate sprouting of afferent (vGluT1^+^) fibres supraspinal injury and neurotrophin-3 treatment [1].

**Figure 1:**
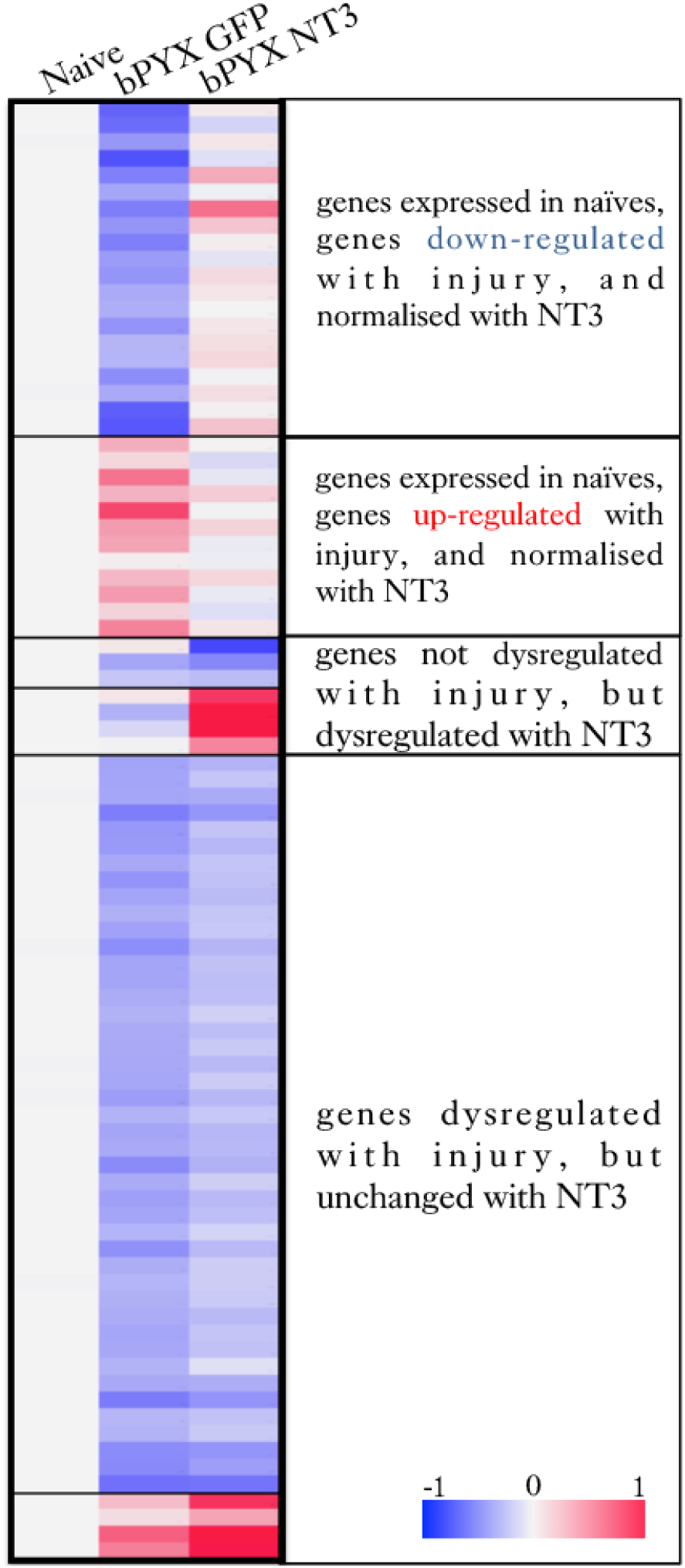
After bilateral transection of the corticospinal tract in the pyramids (bPYX), rats received unilateral intramuscular injections of either AAV1-CMV-NT3 or AAV1-CMV-GFP. Naïve rats were unoperated. Cervical 6-8 DRGs were removed 10 weeks after surgery. RNAseq of poly(A) RNA showed changes in gene expression between these three groups. Lists of poly(A) RNAs for these comparisons are provided in Supplementary Table 6, Supplementary Table 7 and Supplementary Table 8.

**Figure 2:**
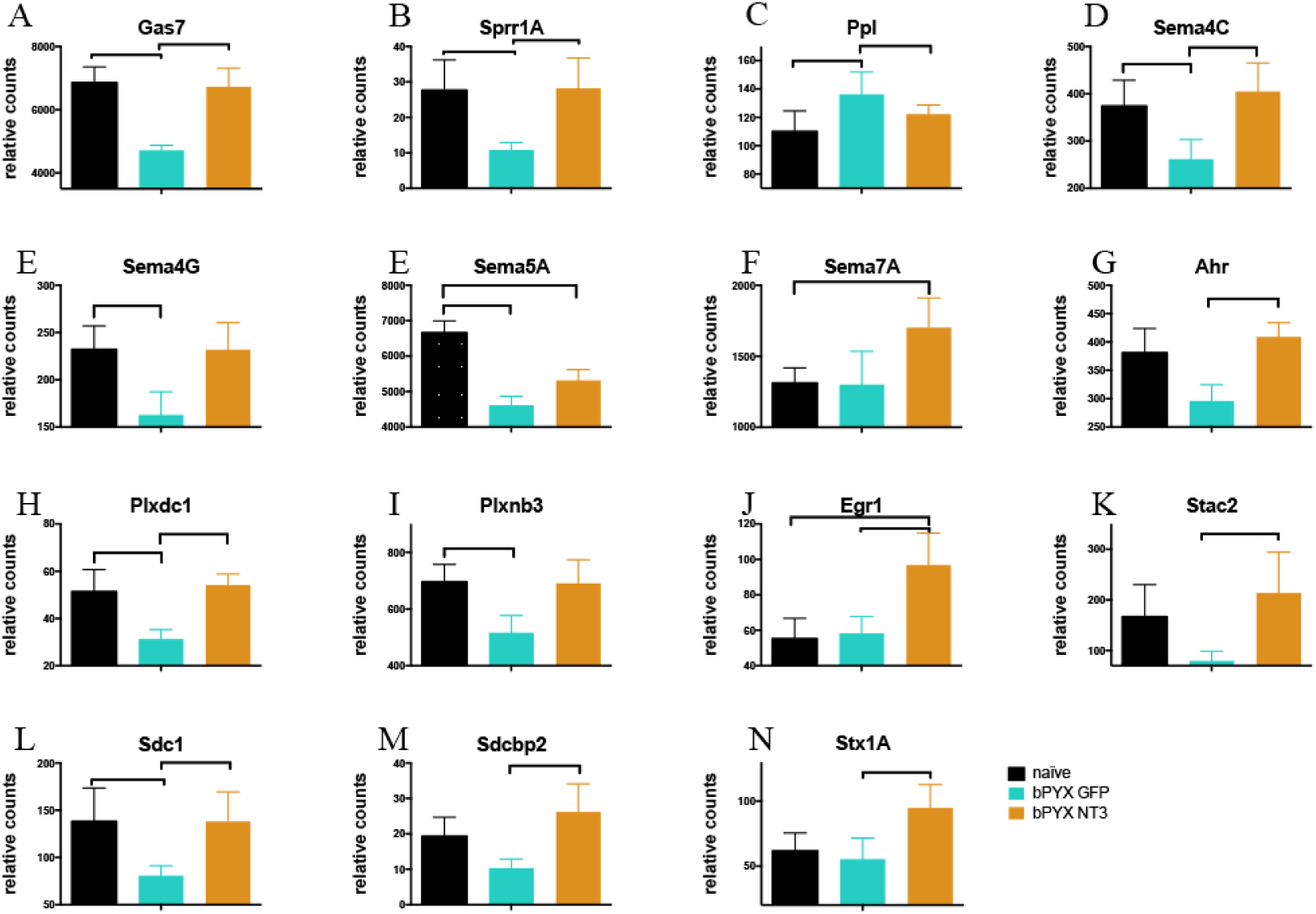
Neurotrophin-3 completely or partially normalizes the levels of many genes in DRG dysregulated by bilateral CST injury. **(A)** - **(N)** RNAseq expression values for selected genes, relative counts. **(A)** - **(C)** *Gas7 Sprr1A* and *Ppl* are important for cytoskeletal reorganization. **(D)** - **(F)** *Sema4C Sema4G Sema5A* and *Sema7A* are transmembrane Semaphorins. **(G)** *Ahr* is a transcription factor regulating Sema4C and Sema7a expression. **(H)** - **(I)** *Plxdc1* and *Plxbn3* belong to the group of plexins, which are binding partners for Semaphorins. **(J)** *Egr1* is a regeneration-associated gene thought to be important for synapse formation. **(K)** - **(N)** *Stac2, Sdc1 Sdcbp2* and *Stx1A* are synapse associated proteins. *n.b.*, y-axes are not always shown from zero upwards. Means ± SD. Brackets indicate p<0.05.

**Figure 3:**
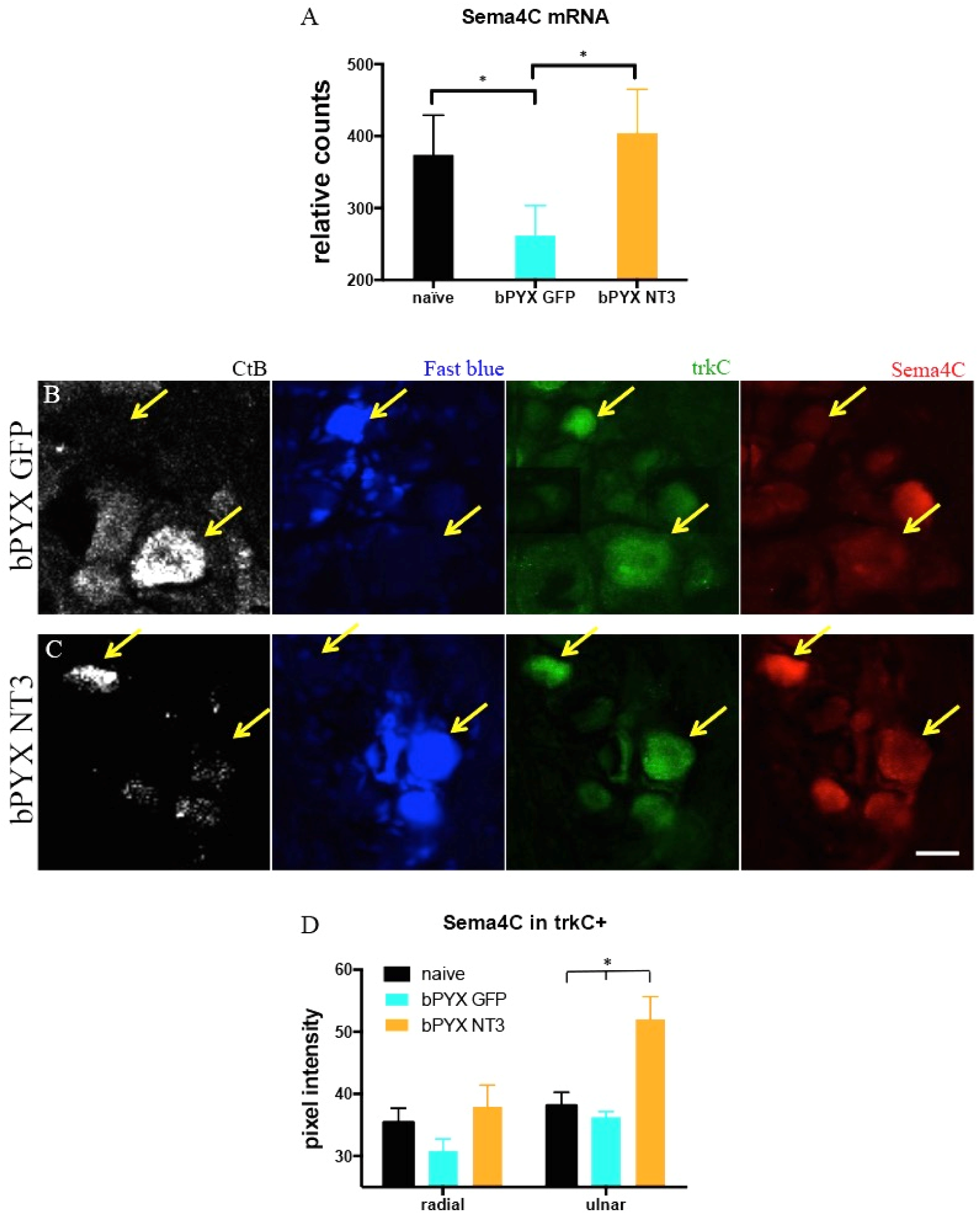
AAV1-CMV-NT3 restores the expression of the axon guidance molecule Sema 4C in cervical DRG neurons. **(A)** RNAseq showed that *Sema4C mRNA*, a transmembrane axon guidance molecule, was downregulated in bPYX+GFP animals. NT3 normalized Sema4C mRNA levels. **(B-C)** Ulnar and radial dorsal root ganglion neurons were traced with CTb and Fast blue respectively. They were immunolabelled with antibodies against TrkC and Sema4C. Yellow arrows indicated CTb^+^ TrkC^+^ and Fast blue^+^ TrkC^+^ example neurons, which were analysed for Sema4C staining intensity. **(D)** Pixel intensity was increased in ulnar nerve TrkC^+^ dorsal root ganglia neurons after neurotrophin-3 treatment (two-way ANOVA, group F=11.5 p<0.001; ulnar nerve DRG neurons bPYX+NT3 vs naive/bPYX+GFP, p-values<0.01). Means ± SDs.

To identify genes that might be regulated in TrkC+ neurons by NT3 in our experiment, we took advantage of a publicly available microarray dataset (GSE38074) containing information about expression of mRNAs (but lacking information about small RNAs) in fluorescence-activated cell sorted TrkC^+^ neurons from the postnatal day 0 DRG either of mice with increased muscular neurotrophin-3 (mlcNT3 versus wildtype control) or of mice lacking neurotrophin-3 (on a Bax knockout background to prevent cell death during development; NT3KO BaxKO versus BaxKO) [7]. We sought genes whose levels correlated positively with NT3 levels and discovered five genes (Spry4, Rasgef1c, Rtn4ip1, Plk2 and Vgf; Table 2) whose levels were higher in our main comparison (bPYX+NT3 vs bPYX+GFP) and also higher in mlcNT3 vs wildtype and also lower in NT3KO BaxKO vs BaxKO (p-values<0.05; see Table 2 for fold changes). Regulation of several of these genes in DRG by neurotrophin-3 is consistent with the literature. For example, Vgf (*n.b.*, not acronymic and not Vegf) is a secreted peptide known to be expressed in sensory ganglia and whose expression is upregulated by neurotrophins [12]. We did not find any genes whose levels correlated negatively with NT3 levels in all three comparisons (i.e., were lower in bPYX+NT3 vs bPYX+GFP and lower in mlcNT3 vs wildtype and higher in NT3KO BaxKO vs BaxKO). Future work will show whether these genes are expressed in adult TrkC+ neurons and whether they are upregulated by NT3, as our data suggests.

Another gene of interest was Sema4C (Figure 3). Semaphorins are axon guidance molecules, which have been shown to be important for specificity of afferent proprioceptive connectivity within the spinal cord [13]; they can be attractants or repellents for proprioceptive afferents [14]. Sema4C is anchored in the membrane and its binding partner Plexin B2 is expressed by spinal motor neurons as well as other spinal neurons (Allen Brain Atlas; adult mouse spinal cord). RNAseq showed that *Sema4C* was down-regulated in cervical DRG after corticospinal tract injury (Figure 3A), which (if it mediates repulsion on contact) might result in proprioceptive vGluT1+ afferents stabilising more contacts onto motor neurons; this might partly explain the changes in proprioceptive afferent connectivity we observed using tracing and polysynaptic reflex testing [1]. RNAseq showed that intramuscular neurotrophin-3 treatment normalized the expression of *Sema4C mRNA* in DRG (Figure 3A; Figure 3).

Next, we evaluated with immunolabeling whether neurotrophin-3 treatment regulates Sema4C expression specifically in TrkC^+^ afferent neurons, which include proprioceptive afferents. We distinguished between ulnar and radial afferents by retrograde tracing with Cholera toxin beta (Ctb) subunit and Fast blue respectively (Figure 3B, C). The ulnar nerve contains afferents from forelimb muscles that were treated with intramuscular AAV-NT3, whereas the radial nerve contains afferents from non-injected muscle groups. We found that TrkC^+^ neurons with ulnar afferents had increased Sema4C expression after intramuscular neurotrophin-3 treatment, but not the cell bodies with afferents from the radial nerve (Figure 3B-D).

These changes may underlie the anatomical and neurophysiological changes in proprioceptive circuits which occur after corticospinal tract injury and which are normalized by neurotrophin-3 treatment [1].

This dataset will enable us and others to identify genes involved in changes in sensory axon anatomy and function after CNS injury.

## Experimental Design, Materials and Methods

As described previously, bilateral pyramidotomy was performed in rats and AAV1-CMV-NT3 or AAV1-CMV-EGFP was injected into forelimb flexor muscles unilaterally [1]. These rats underwent behavioural testing, neurophysiological assessment and nerve tracing as described elsewhere [1]. Cervical DRG 6 to 8 were dissected from unfixed tissues and snap-frozen in liquid nitrogen. We isolated RNA from homogenates of DRGs to assess overall gene expression in neurons and glia.

### Total RNA extraction for Sequencing

Total RNA for sequencing was extracted from C6-C8 DRGs which were pooled unilaterally from the treated side per animal (and not between animals). Tissue was homogenized in QIAZOL (Qiagen, 79306) and nucleic acids were separated in Phase Locked Gel columns (5Prime, 230 2830). 1.5 volume of 100% ethanol was added to the aqueous phase and transferred to filtered spin columns from the extraction kit for poly A RNA and small RNAs (miRNeasy kit, Qiagen, 217004) and we proceeded according to manufacturer’s instructions. Samples were also DNase I treated with double volume of the recommended amount (Qiagen, 79254). We estimated the total RNA quality and quantity by spectrophotometry (NanoDrop, ND-1000) and measured the RNA integrity numbers (RINs) (Agilent RNA 6000 Nano Reagents Part I and Agilent 2100 Bioanalyzer). The RIN for each sample was greater than 7.6, with an average of 8 (Supplementary Table 1).

### Poly A RNA sequencing and analysis

The samples were subjected to poly(A) enrichment with oligo-dT beads (Illumina) and the library was prepared using the TruSeq Stranded prep kit (Illumina). Samples were run as a barcoded multiplex over 4 lanes on the Illumina HiSeq 2500 platform using the Rapid SBS kit v2 (50bp read length, paired end). Illumina universal paired end adapters were used:

5’ P-<u>**GATCGGAAGAGC**</u>GGTTCAGCAGGAATGCCGAG

5’ ACACTCTTTCCCTACACGAC<u>**GCTCTTCCGATC**</u>T

The success of the sequencing run and quality of the raw data were assessed by a range of metrics using custom scripts. The 50bp paired-end reads were aligned to the Rattus norvegicus reference genome (Rnor_6.0) using TopHat2 with default parameters (except for setting mate-inner-dist=100 and mate-std-dev=50). Just over 30 million read-pairs per sample were obtained on average (mean ± SD, 32.1±4.4 million) and around 94% of these could be aligned to the reference genome. Duplicate reads were identified using Picard Tools MarkDuplicates (Picard Tools by The Broad Institute) and the highest quality read at each position was retained. Reads mapping to each gene feature were counted using HTseq to create a raw gene count table for 15,075 RefSeq annotated genes with at least one mapped read (Supplementary Table 2). Due to a relatively large number of reads annotated as ‘no feature’ (i.e. mapping to intronic or intergenic regions) and exclusion of duplicate reads and those mapping to multiple locations, the total number of read-pairs mapped to gene features per sample was 12.8±1.8 million (mean ± SD). Further quality control plots and exploratory analyses including principal component analysis (PCA) were performed to assess the overall behaviour of the dataset. Although minor outliers were observed, all samples were included for subsequent analysis for mRNA expression profiles. A detection filter requiring >10 reads on average in at least 6 samples was applied and 11,528 genes were considered expressed and retained for differential expression analysis using the EdgeR package comparing and creating the following three data sets: bPYX+GFP vs naive, bPYX+NT3 vs bPYX+GFP, bPYX+NT3 vs naive. Lists of differentially regulated poly(A) RNAs were based on p<0.05; changes in expression level (log_2_ fold change) for all poly(A) RNAs (i.e., whether significantly regulated or not) for all three comparisons is provided in Supplementary Table 4.

### Small RNA sequencing and analysis

Libraries were prepared using the Small Library Prep Set for Illumina (NEBNext, multiplex compatible; NEB) using custom index primers. Samples were run as a barcoded multiplex over 4 lanes on the Illumina HiSeq 2500 platform using the Rapid SBS kit v2 (50bp read length, single end).

miRNA data was mapped to RNor_5.0, in the absence of a miRNA resource for rn6 at that time. The total number of reads was around 13 million (Mean: 13.5M, SD: 1.7M). Of those, 96.7% mapped to the target regions and 93.5% mapped to miRNA features. For miRNA sequencing, the analysis pipeline included adapter trimming prior to mapping, and used the short-read aligner Bowtie2. The count table (Supplementary Table 3) was generated with HTSeq-count. RNA sequencing analyses including QC and exclusion of samples from analysis was performed by an impartial third party bioinformatician: one outlier (rat #41) was identified in the QC plots and excluded for further analysis. A detection filter requiring >10 reads on average in at least 6 samples was applied and 363 small RNAs were considered expressed and retained for differential expression analysis using the EdgeR package comparing and creating following data sets: bPYX+GFP vs naive, bPYX+NT3 vs bPYX+GFP, bPYX+NT3 vs naive. Lists of differentially regulated small RNAs were based on p<0.05. Changes in expression level (log_2_ fold change) for all small RNAs (*i.e.*, whether significantly regulated or not) for all three comparisons is provided in Supplementary Table 5.

### Bioinformatic analyses

Next, three lists of poly(A) RNAs were created that were differentially regulated in two of our comparisons. This revealed genes that were 1) dysregulated after injury and then normalized with NT3 treatment, 2) dysregulated after injury and not normalised with NT3 treatment, 3) not dysregulated after injury, but dysregulated with NT3 treatment (Supplementary Table 6, Supplementary Table 7, Supplementary Table 8).

To identify genes that might be regulated in TrkC+ neurons by NT3 in our experiment, we next took advantage of a publicly available microarray dataset (GSE38074) containing information about expression of mRNAs (but lacking information about small RNAs) in fluorescence-activated cell sorted TrkC^+^ neurons from the postnatal day 0 DRG either of mice with increased muscular NT-3 (mlcNT3 versus wildtype control; n=2/group) or of mice lacking NT-3 (on a Bax knockout background to prevent cell death during development; NT3KO BaxKO versus BaxKO; n=2/group) [7]. We sought genes whose levels correlated positively with NT3 levels in these two comparisons and in bPYX+NT3 versus bPYX+GFP.

### Immunofluorescence staining

Tissues were obtained from a previous study [1]. Immediately after performing nerve neurophysiology, rats were transcardially perfused with PBS pH 7.4 and tissues were dissected rapidly. Tissue for sectioning on the cryostat was post-fixed by immersion in 4% paraformaldehyde in PBS pH 7.4 overnight and cryoprotected in 30% sucrose in PBS pH 7.4. Tissue was frozen and embedded in O.C.T., DRGs were sectioned transversely at 10 μm thickness respectively and directly mounted onto glass slides. Immunofluorescence staining was performed [1] using antibodies shown below (Table 1). All antibodies were verified by their manufacturer. The Sema4C antibody is raised against a peptide mapping with a C-terminal cytoplasmic domain of Sema4C human origin.

**Table 1:**
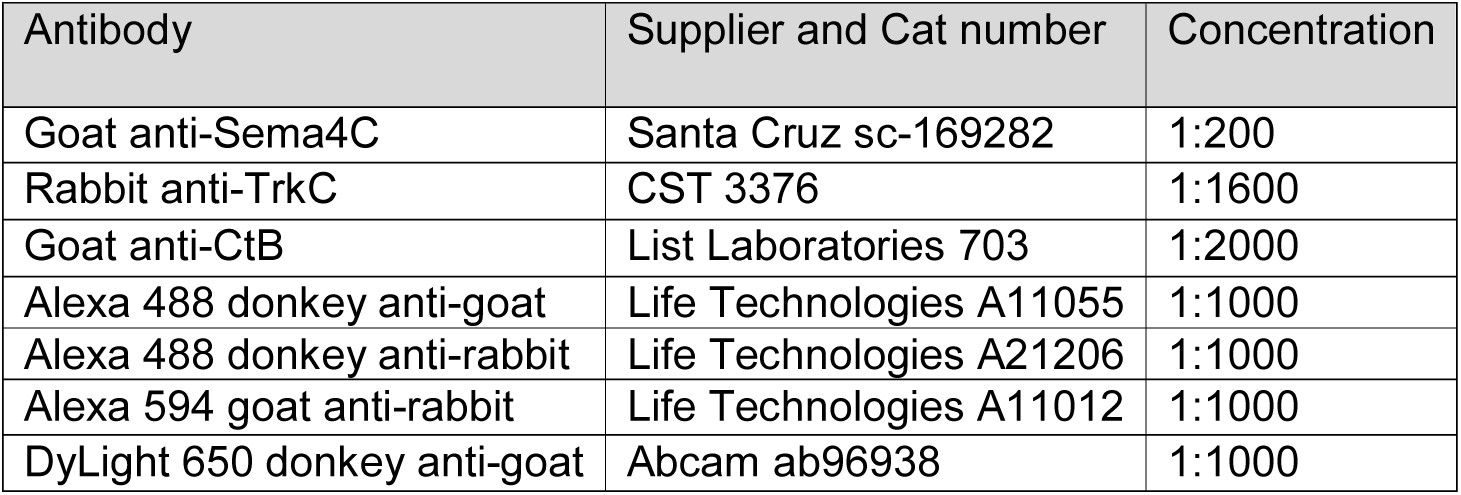
Table showing primary and secondary antibodies

| Antibody | Supplier and Cat number | Concentration |
| --- | --- | --- |
| Goat anti-Sema4C | Santa Cruz sc-169282 | 1:200 |
| Rabbit anti-TrkC | CST 3376 | 1:1600 |
| Goat anti-CtB | List Laboratories 703 | 1:2000 |
| Alexa 488 donkey anti-goat | Life Technologies A11055 | 1:1000 |
| Alexa 488 donkey anti-rabbit | Life Technologies A21206 | 1:1000 |
| Alexa 594 goat anti-rabbit | Life Technologies A11012 | 1:1000 |
| DyLight 650 donkey anti-goat | Abcam ab96938 | 1:1000 |

**Table 2:** The levels of five mRNAs correlated with higher NT3 in all three key comparisons: bPYX+NT3 v bPYX+GFP, mlcNT3 vs wildtype and NT3KO BaxKO v BaxKO (p<0.05).

|  | Log<sub>2</sub> fold change<br>in bPYX+NT3 vs<br>bPYX+GFP | Log<sub>2</sub> fold change<br>in mlcNT3 vs<br>wildtype | Log<sub>2</sub> fold change<br>in NT3KO BaxKO<br>versus BaxKO |
| --- | --- | --- | --- |
| Spry4 | 0.4 | 2.2 | -1.7 |
| Rasgef1c | 0.3 | 0.4 | -0.6 |
| Rtn4ip1 | 0.3 | 0.4 | -0.8 |
| Plk2 | 0.2 | 0.6 | -1.2 |
| Vgf | 0.5 | 1.2 | -2.5 |

### Image Analysis

Image analysis was performed with Image J or Zen Imaging software. Pixel intensity of Sema4C staining was analysed in dorsal root ganglia neurons, which contained afferents from ulnar (CTb positive) and radial (Fast blue positive) nerve. The pixel intensity of the cytoplasm of TrkC^+^ neurons was measured. 8 neurons per section and 3 sections per animal were analysed. The mean number per animals was calculated. The mean was averaged within the groups.

## Acknowledgements

We thank Dr Ana Antunes-Martins for advice regarding preparation of RNA samples. We thank the High-Throughput Genomics Group at the Wellcome Trust Centre for Human Genetics (funded by Wellcome Trust grant reference 090532/Z/09/Z) for the generation of the Sequencing data. We especially thank Drs. Simon Engledow, Helen Lockstone and Benjamin Wright (Oxford Genomics Centre, Wellcome Trust Centre for Human Genetics) for RNA sequencing and analysis. The research leading to these results has received funding from the International Spinal Research Trust’s Nathalie Rose Barr Studentship (NRB106), a Serendipity grant from the Dunhill Medical Trust (SA21/0521), The Rosetrees Trust, the European Research Council under the European Union’s Seventh Framework Programme (FP/2007-2013) / ERC Grant Agreement n. 309731, and King’s College London Graduate Teaching Assistant Program.

## Supplementary Tables

**Supplementary Table 1:** Table showing sample identification numbers submitted for RNA sequencing together with sample concentration (ng/ul), ratio of absorbance readings at 260nm/280nm and at 260nm/230nm, RIN score. Group mp1 = naïve; Group mp2 = bPYX+GFP; Group mp3 = bPYX+NT3.

| Sample Name | RNA Conc. | Units | Abs 260/280 | UDF/260/230 | Group | RIN score |
| --- | --- | --- | --- | --- | --- | --- |
| 2 | 196.0 | ng/ul | 2.09 | 2.25 | mp3 | 8.3 |
| 5 | 191.6 | ng/ul | 2.08 | 1.93 | mp2 | 8.3 |
| 11 | 189.5 | ng/ul | 2.06 | 2 | mp3 | 7.9 |
| 15 | 177.8 | ng/ul | 2.14 | 2.12 | mp2 | 8 |
| 17 | 160.7 | ng/ul | 2.1 | 2.46 | mp3 | 7.8 |
| 18 | 154.1 | ng/ul | 2.06 | 1.61 | mp2 | 7.6 |
| 21 | 166.2 | ng/ul | 2.11 | 2.3 | mp2 | 8 |
| 23 | 192.5 | ng/ul | 2.09 | 2.34 | mp3 | 7.6 |
| 39 | 180.2 | ng/ul | 2.07 | 2.25 | mp3 | 8 |
| 40 | 163.3 | ng/ul | 2.13 | 1.98 | mp3 | 7.9 |
| 41 | 172.2 | ng/ul | 2.1 | 2.36 | mp1 | 8.2 |
| 42 | 170.5 | ng/ul | 2.07 | 2.27 | mp1 | 7.9 |
| 43 | 181.6 | ng/ul | 2.04 | 2.03 | mp1 | 8.1 |
| 44 | 201.0 | ng/ul | 2.02 | 1.98 | mp1 | 7.8 |
| 45 | 230.4 | ng/ul | 2.08 | 2.35 | mp1 | 8.3 |
| 49 | 149.9 | ng/ul | 2.05 | 2.33 | mp1 | 8.4 |
| 52 | 148.0 | ng/ul | 2.1 | 2.41 | mp2 | 8.1 |
| 57 | 169.1 | ng/ul | 2.13 | 2.37 | mp2 | 8.2 |

Supplementary Table 2: Excel file containing count data for 15,075 poly(A) RNAs in cervical sensory ganglia with at least one mapped read. Sample names are as shown in Supplementary Table 1. Group means and standard deviations are shown.

Supplementary Table 3: Excel file showing counts per sample for 753 small RNAs. Sample names are as shown in Supplementary Table 1. Group means and standard deviations are shown for all samples and also when outlier sample #41 was omitted.

Supplementary Table 4: Excel file containing data for poly(A) RNAs in cervical sensory ganglia. Three comparisons are shown (one per worksheet); bPYX+GFP *versus* naïve; bPYX+NT3 *versus* naïve; bPYX+NT3 *versus* bPYX+GFP. Each list shows all poly(A) RNAs above the cut-off threshold, whether or not significantly regulated. Log_2_ fold change is positive when the first named group in that worksheet has higher expression level than the second named group in that worksheet (e.g., bPYX+NT3 *versus* bPYX+GFP).

Supplementary Table 5: Excel file containing data for small RNAs in cervical sensory ganglia. Three comparisons are shown (one per worksheet); bPYX+GFP *versus* naïve; bPYX+NT3 *versus* naïve; bPYX+ NT3 *versus* bPYX+GFP. Each list shows all small RNAs above the cut-off threshold, whether or not significantly regulated. Table shows miRNA name, log_2_ fold change, log CPM, LR, PValue and FDR. Log_2_ fold change is positive when the first named group in that worksheet has higher expression level than the second named group in that worksheet (e.g., bPYX+NT3 *versus* bPYX+GFP). Sample #41 was omitted as it was an outlier.

**Supplementary Table 6:**
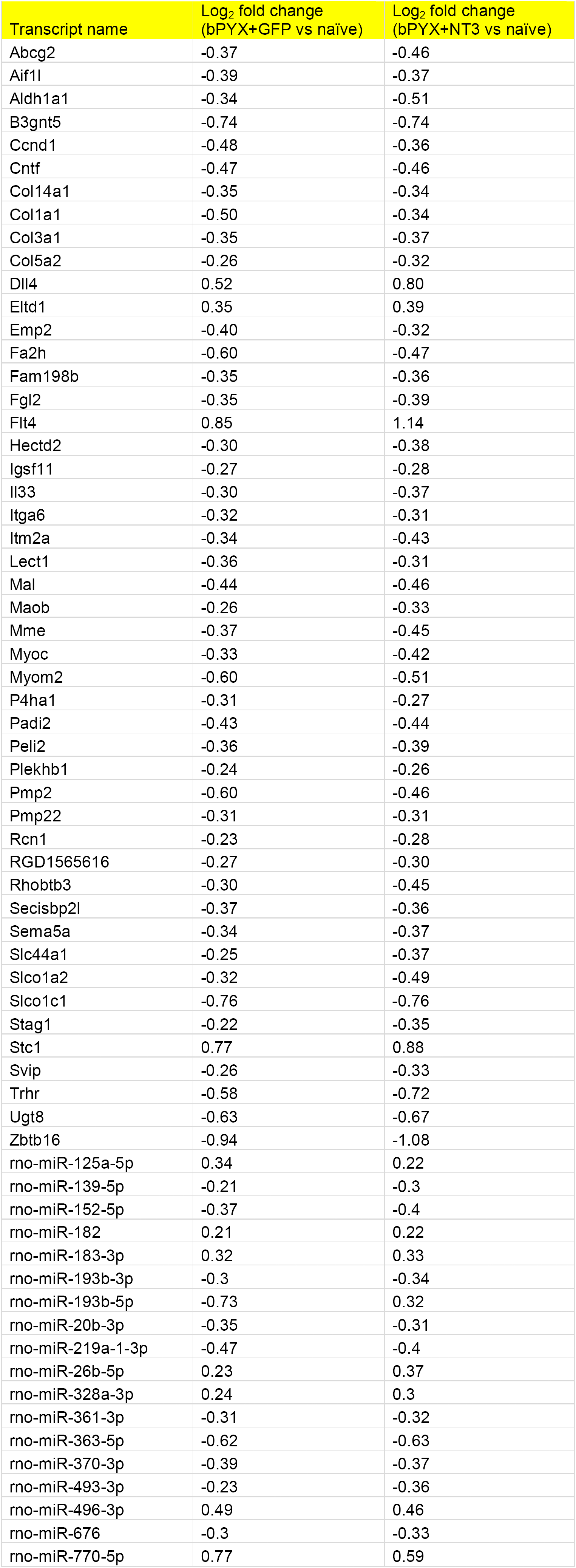
Sequencing identified 48 mRNAs and 18 small RNAs in cervical sensory ganglia whose expression levels were modified by bPYX+GFP *versus* the naïve group (p<0.05) and not by bPYX+NT3 *versus* bPYX+GFP (p>0.05).

**Supplementary Table 7:** Sequencing identified 32 mRNAs and 9 small RNAs in cervical sensory ganglia whose expression levels were modified by pyramidotomy (bPYX+GFP versus naïve, p<0.05) and reversed by intramuscular NT3 treatment relative to pyramidotomy plus GFP (bPYX+NT3 versus bPYX+GFP, p<0.05).

| Transcript name | Log<sub>2</sub> fold change<br>(bPYX+GFP vs naïve) | log<sub>2</sub> fold change<br>(bPYX+NT3 vs bPYX+GFP) |
| --- | --- | --- |
| Abca4 | 0.55 | -0.57 |
| Arg1 | -1.01 | 1.10 |
| Aurkb | -0.94 | 0.73 |
| Cdkn1a | -0.45 | 0.49 |
| Cldn11 | 0.36 | -0.53 |
| Crisp1 | -1.28 | 1.20 |
| Cxcl14 | -0.71 | 1.01 |
| Fmo3 | 0.78 | -0.79 |
| Frzb | 0.54 | -0.36 |
| Gas7 | -0.35 | 0.26 |
| Gfap | -0.70 | 1.26 |
| Gpnmb | 0.86 | -0.89 |
| Id1 | 0.60 | -0.45 |
| Loxl4 | -0.51 | 0.72 |
| Ly6g6e | -0.74 | 0.70 |
| Myo10 | -0.45 | 0.33 |
| Myof | 0.55 | -0.62 |
| Net1 | 0.23 | -0.28 |
| Nfil3 | -0.51 | 0.60 |
| Pdia5 | -0.32 | 0.35 |
| Plk2 | -0.29 | 0.25 |
| Plxdc1 | -0.49 | 0.56 |
| Ppl | 0.49 | -0.36 |
| Rasa3 | -0.26 | 0.31 |
| Rcan1 | -0.23 | 0.34 |
| Sdc1 | -0.53 | 0.52 |
| Sema4c | -0.33 | 0.40 |
| Sprr1a | -1.11 | 1.13 |
| Tgm1 | -1.21 | 1.45 |
| Upk1b | 0.64 | -0.68 |
| Zic2 | 0.38 | -0.51 |
| Zic5 | 0.73 | -0.66 |
| rno-let-7b-3p | 0.25 | -0.3 |
| rno-miR-145-5p | -0.45 | 0.35 |
| rno-miR-146b-5p | -0.6 | 0.36 |
| rno-miR-148a-3p | 0.77 | -0.66 |
| rno-miR-148b-3p | 0.63 | -0.61 |
| rno-miR-30d-3p | -0.4 | 0.31 |
| rno-miR-30e-3p | 0.45 | -0.26 |
| rno-miR-344b-1-3p | -0.53 | 0.34 |
| rno-miR-490-3p | -0.3 | 0.26 |

**Supplementary Table 8:** Sequencing identified seven mRNAs and seven small RNAs in cervical sensory ganglia whose expression levels were modified by bPYX+NT3 *versus* bPYX+GFP (p<0.05) or bPYX+NT3 *versus* naïve (p<0.05) and not in bPYX+GFP versus naïve (p>0.05).

| Transcript name | Log<sub>2</sub> fold change<br>(bPYX+NT3 vs bPYX+GFP) | Log<sub>2</sub> fold change<br>(bPYX+NT3 vs naïve) |
| --- | --- | --- |
| Adipoq | -2.11 | -1.85 |
| Cpa3 | -0.53 | -0.88 |
| Egr1 | 0.49 | 0.76 |
| Mall | 1.15 | 0.90 |
| Map3k14 | 0.86 | 0.87 |
| Ncan | 0.31 | 0.48 |
| Prex2 | -0.28 | -0.38 |
| rno-miR-181c-5p | 0.53 | 0.5 |
| rno-miR-582-5p | 0.71 | 0.57 |
| rno-miR-193a-3p | 1.66 | 1.76 |
| rno-miR-137-3p | 0.4 | 0.71 |
| rno-miR-205 | 0.44 | 0.45 |
| rno-miR-22-3p | 0.45 | 0.36 |
| rno-miR-29a-5p | 0.64 | 0.55 |

